# Brd2 is dispensable for genome compartmentalization and replication timing

**DOI:** 10.1101/2023.11.17.567572

**Authors:** Laura Hinojosa-Gonzalez, Jesse L. Turner, Takayo Sasaki, Ferhat Ay, David M. Gilbert

## Abstract

Replication Timing (RT) refers to the temporal order in which the genome is replicated during S phase. Early replicating regions correlate with the transcriptionally active, accessible euchromatin (A) compartment, while late replicating regions correlate with the heterochromatin (B) compartment and repressive histone marks. Previously, widespread A/B genome compartmentalization changes were reported following Brd2 depletion. Since RT and A/B compartmentalization are two of the most highly correlated chromosome properties, we evaluated the effects of Brd2 depletion on RT. We performed E/L Repli-Seq following Brd2 depletion in the previously described Brd2 conditional degron cell line and found no significant alterations in RT after Brd2 KD. This finding prompted us to re-analyze the Micro-C data from the previous publication. We report that we were unable to detect any compartmentalization changes in Brd2 depleted cells compared to DMSO control using the same data. Taken together, our findings demonstrate that Brd2 depletion alone does not affect A/B compartmentalization or RT in mouse embryonic stem cells.

## INTRODUCTION

The genome is replicated in a defined and tightly regulated temporal order known as the replication timing (RT) program. RT programs are evolutionarily conserved, cell-type specific, and particularly resistant to a wide range of protein depletions, knockdowns, knockouts, and inhibitors, indicating robust control (Dileep et al. 2015; Marchal et al. 2019). RT is essential for proper epigenome maintenance (Klein et al. 2021) and irregularities in the RT program are associated with genetic instability and numerous diseases and cancers (Vouzas and Gilbert 2023). Despite these findings, the precise mechanisms that control RT are poorly understood. One longstanding observation is the close relationship between RT and genome compartmentalization. Early replication takes place in the active interior of the nucleus and late replication in the inactive heterochromatin often found at the nuclear and nucleolar periphery (O’Keefe et al. 1992; Zink 2006). The first molecular measurements of chromatin interactions by Hi-C also defined open and closed chromatin compartments termed A and B, respectively. Comparison of these two properties revealed an exceptionally strong correlation with RT, demonstrating that Hi-C A/B compartments correspond to the cytologically visualized early and late replicating spatial compartments of chromatin in the nucleus (Ryba et al. 2010; Yaffe et al. 2010).

A recent study (Xie et al. 2022) reported that the bromodomain and extraterminal (BET) family scaffold protein Brd2 is essential for compartmentalization of the accessible genome in mouse embryonic stem cells (mESC). Since A/B compartmentalization and RT are highly correlated, we hypothesized that Brd2 KD could identify a rare genetic perturbation capable of causing widespread perturbations in RT. Here, we performed RepliSeq after 24 hours of Brd2 depletion in the same previously described Brd2 dTAG cell line. Surprisingly, we did not find any statistically significant changes in RT. This led us to revisit the Xie et. al. compartment analysis.

The standard approach for defining compartments from contact matrices is to perform principal component analysis (PCA) on the intra-chromosomal correlation matrix for each chromosome. However, A/B compartment analysis is not straightforward because the first principal component (the eigenvector that corresponds to the largest eigenvalue) is not always the one that captures chromatin compartments (Lieberman-Aiden et al. 2009). It is important to utilize GC content, gene expression, or other genomic features to choose the correct PC and orient the scores properly so positive values represent A compartments and negative values represent B compartments. Our re-analysis of the published Micro-C data revealed that Brd2 KD does not impact A/B compartmentalization. We conclude that A/B compartmentalization and RT are robust to Brd2 KD and remain tightly correlated in both control and Brd2 depletion conditions.

## RESULTS AND DISCUSSION

### Brd2 depletion does not affect Replication Timing

We were generously gifted the Brd2-FKBP^F36V^ mESC cell line (Xie et al. 2022) in which both copies of an endogenous Brd2 gene were tagged with human FKBP12 allele FKBP^F36V^ (degron), to achieve rapid and reversible degradation of Brd2 upon the addition of dTAG-13 (dTAG) (Nabet et al. 2018). The RT program was evaluated in the presence and absence of dTAG for 24 hours using E/L Repli-seq (Marchal et al. 2018) after confirming Brd2 depletion with dTAG **(Supplementary Fig 1)**. Genome-wide analyses could identify no significant differences between the Brd2 degron (dTAG) and control (DMSO) **(Fig. 1A-B; Supplementary Fig 2)**. Since large localized B to A switches were reported in Chr11 and 17 following Brd2 depletion (Figs. 3B and ED5B in Xie et. al.), we specifically examined RT profiles in those same regions **(Fig. 1C-D)** but again did not observe any RT shifts. Initially, given the claims of the prior report, this suggested a surprising genome-wide uncoupling of the RT program and A/B compartments.

**Fig 1.**
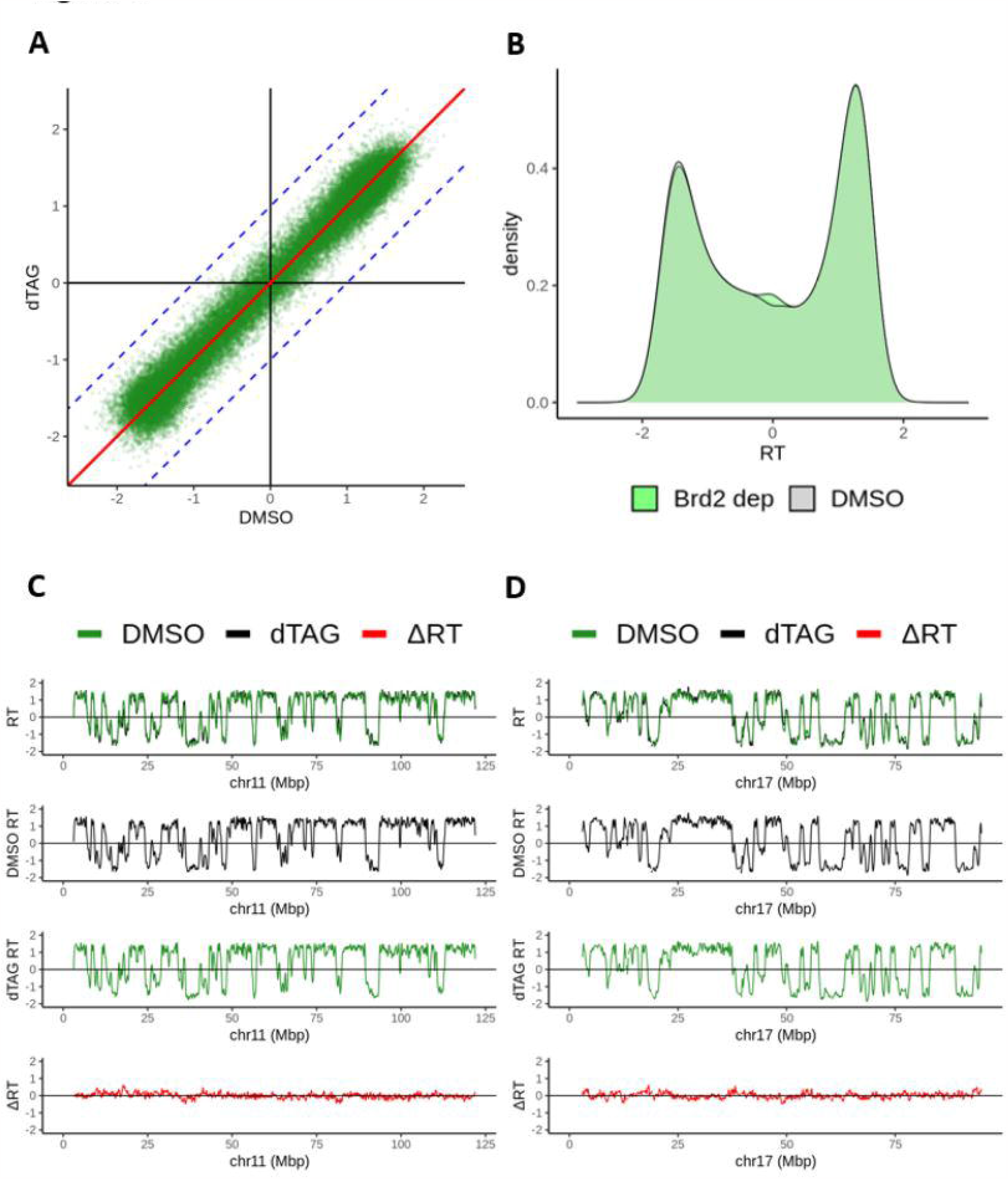
Impact of Brd2 depletion on replication timing (RT). **A** Genome-wide RT values (50kb resolution) between control (DMSO) and Brd2 depletion (dTAG). **B**. Density distribution of RT values plotted in **(A). C-D**. RT profiles for **(C)** chromosome 11 and **(D)** chromosome 17 of control (DMSO; black), Brd2 depletion (dTAG; green), and ΔRT (dTAG – DMSO; red).

### Brd2 depletion does not affect AB Compartmentalization

Because of the long-standing correlation between RT and A/B compartments, we reinvestigated the compartment analysis of the Xie at al. paper to interpret our perplexing findings. We obtained fastqs from GEO/SRA (accession ID: GSE163729), reprocessed the Micro-C data and built contact matrices (see Methods) for the control and Brd2 depletion conditions. We then employed dcHiC (Chakraborty et al. 2022) for compartment analysis (see Methods). Surprisingly, we found no genome-wide changes in compartment score distribution **(Fig. 2A-B; Supplementary Fig 3)** in contrast to what was reported by Xie et al. in their Figure 3A. As with RT, we also looked at chromosomes 11 and 17 specifically (<u>Figs. 3B and ED5B in Xie et. al.</u>), and could not replicate the reported B to A switches and large-scale shifts either **(Fig. 2C-D)**.

**Fig 2.**
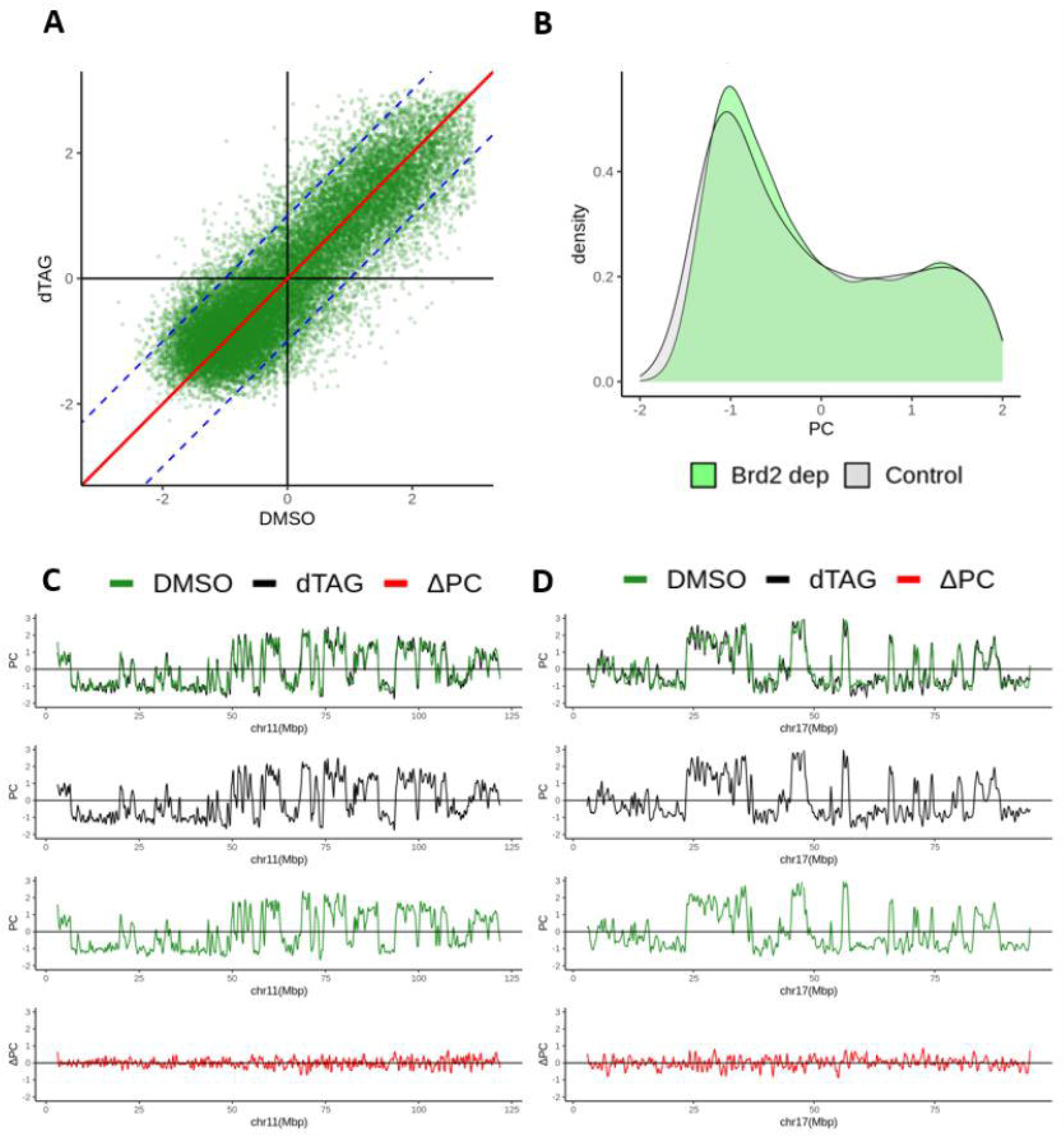
Impact of Brd2 depletion on A/B compartments (re-analysis of Xie et al. data). Genome-wide compartment scores (100kb resolution) between control (DMSO) and Brd2 depletion (dTAG) calculated from Micro-C data (Xie et al. 2022). **B**. Density distribution of compartment values plotted in **(A). C-D**. Compartment score profiles for **(C)** chromosome 11 and **(D)** chromosome 17 of control (DMSO; black), Brd2 depletion (dTAG; green), and ΔPC (dTAG – DMSO; red).

We speculate that the A/B compartmentalization changes reported by Xie et al. are due to errors during compartment analysis, at least for a subset of chromosomes. However, since the original paper did not provide their compartment calls, and they were not available upon request, we could not identify with certainty the source of the discrepancy. In summary, given the lack of alterations in RT after Brd2 knockdown and the general concordance of RT with A/B compartments, combined with our inability to detect compartment changes after Brd2 knockdown, we conclude that Brd2 depletion does not affect A/B compartmentalization or RT in mESCs. We believe that the field remains open to further studies of modulators of genome compartmentalization.

## Author Contributions

L.H-G. performed the computational analysis. J.L.T. and T.S. performed replication timing experiments. F.A. and D.M.G. supervised the work. All authors wrote, reviewed and agreed with the contents of this manuscript.

## DATA AVAILABILITY

All scripts and tables pertaining to this manuscript can be found on GitHub repository: https://github.com/ay-lab/Brd2_analysis. All main analysis can be reproduced through a Jupyter notebook: https://ay-lab.github.io/Brd2_analysis/Plots.html. We provide the 50kb-binned replication timing data. Raw sequencing data will be uploaded to GEO.

## Supplementary Tables

- Supplementary Table 1: Chosen PCs for each chromosome from each sample
- Supplementary Table 2: Replication timing values for control and Brd2 depletion
- Supplementary Table 3: Compartment scores for control and Brd2 depletion

## SUPPLEMENTARY INFORMATION

- Supplementary Figures
  - Supplementary Figure 1: Western blot images
  - Supplementary Figure 2: Per chromosome distributions of replication timing values
  - Supplementary Figure 3: Per chromosome distributions of compartment scores

- Supplementary Methods

## ACKNOWLEDGEMENTS

The Brd2-FKBP^F36V^ mESC cell line was a generous gift from L. Xie. This work was supported by NIH grants R01-GM083337 to DMG and R35-GM128938 to FA.

