## Supplementary Information for "Brd2 is dispensable for genome compartmentalization and replication timing"

**SUPPLEMENTARY FIGURES**

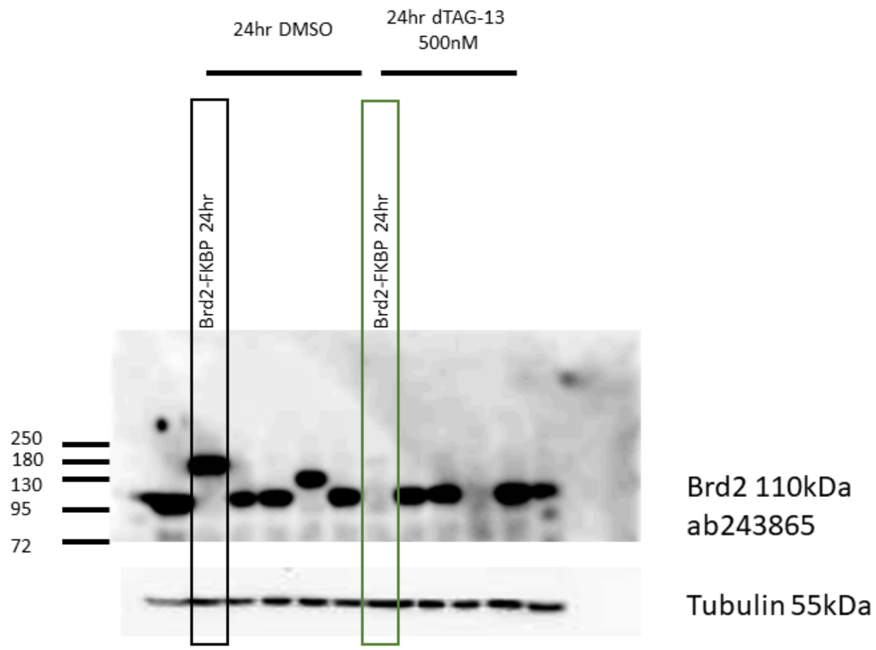


**Supplementary Figure 1.** Validation of Brd2 degradation for E/L Repli-seq by western Blot in Brd2-FKBP.

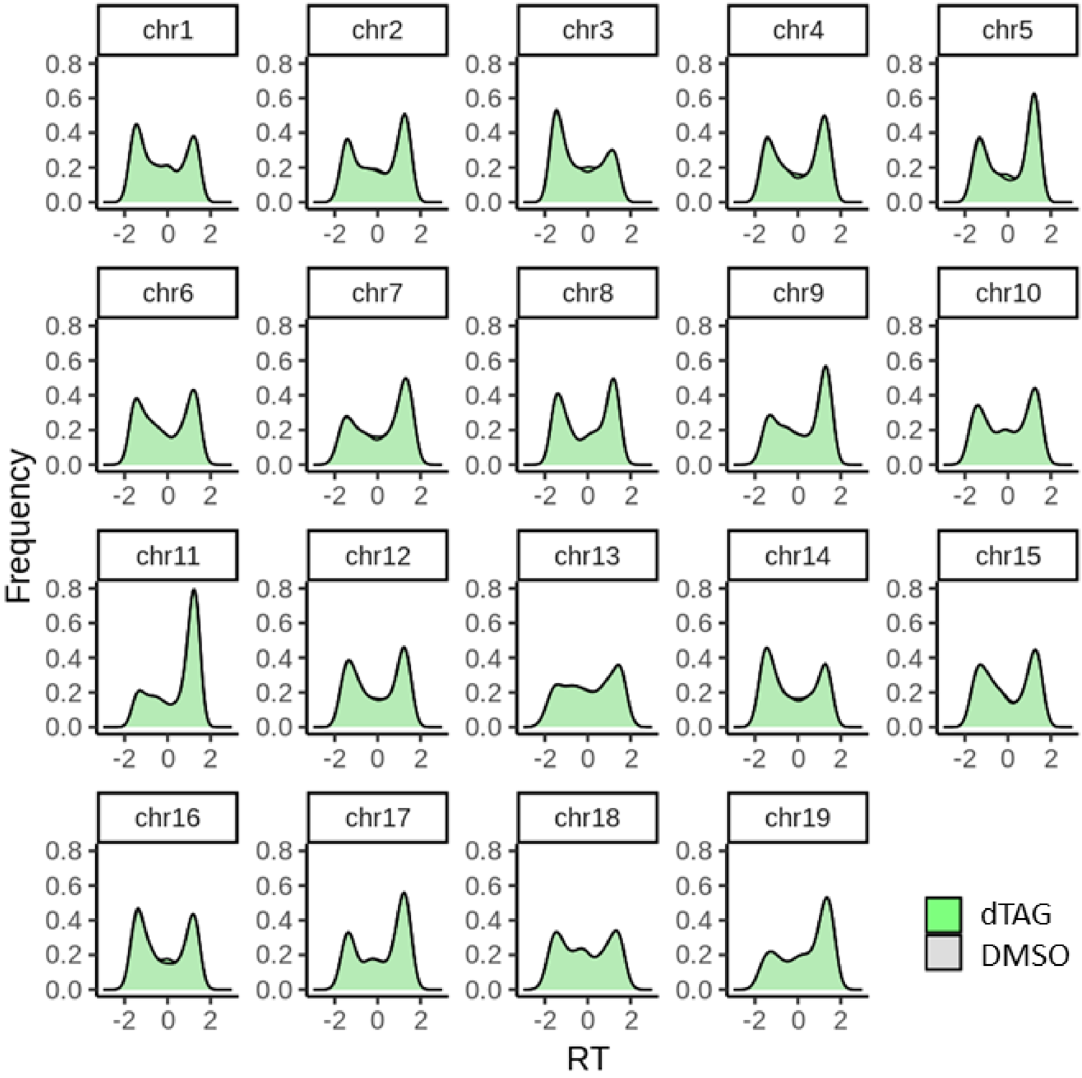


**Supplementary Figure 2.** Per chromosome distribution of replication timing values in Brd2 depletion (dTAG) and control (DMSO).

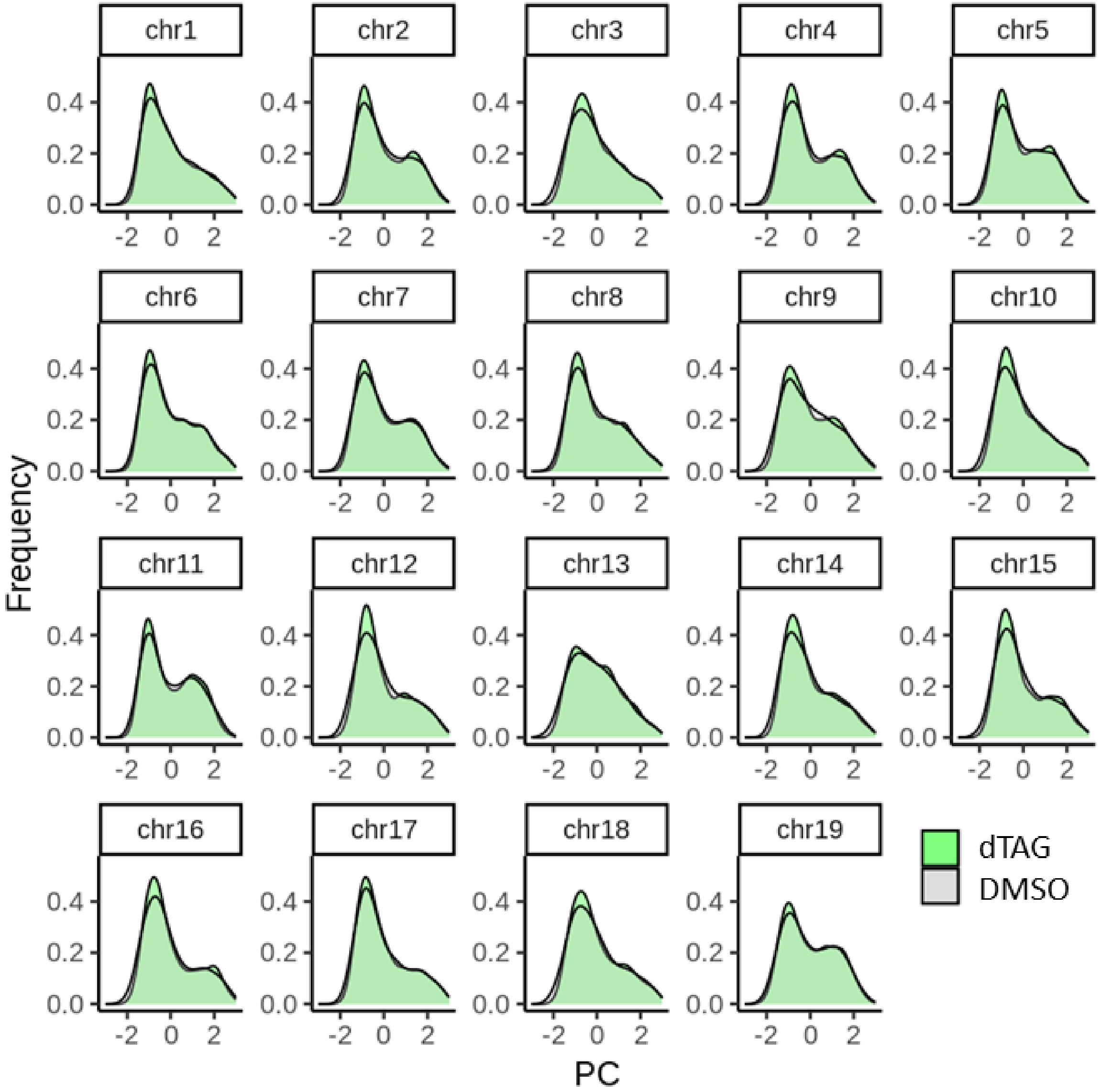


**Supplementary Figure 3.** Per chromosome distribution of PC values in Brd2 depletion (dTAG) and control (DMSO).

### SUPPLEMENTARY METHODS

#### Cell Culture

mESCs were cultured, maintained, and passaged as previously described (Sima et al. 2019; Turner et al. 2023). Briefly, cells were cultured on 0.1% gelatin-coated dishes in serum-free 2i/LIF media. Cell lines obtained from other sources were transitioned into 2i/LIF medium and grown in 2i/LIF for all experiments and manipulations.

#### Degradation (dTAG) Experiments

Cells were cultured in 2i/LIF until reaching proper density. Media was then replaced with either media supplemented with 500nM dTAG-13 in DMSO (MCE, HY-114421) or equivalent volume (0.1%) of DMSO (vehicle). Cells were incubated with drug or DMSO for 22 hours and then labeled for 2 hours with BrdU for Repli-seq (below). Pellets of BrdU-labeled cells were put aside for western blot validation to verify lack of detectable target protein within 2 hours of addition of dTAG-13.

#### Western Blots

200,000 cells equivalent extract per lane was loaded onto mini gel (Bio-Rad Mini Protean III) together with marker (NEB, P7719) and separated at 150V (constant voltage) for 60 minutes on 8% gel for targets. The separated protein was transferred to the PVDF membrane using the Bio-Rad Trans-Blot Turbo Transfer System and Bio-Rad Trans-Blot Turbo RTA Transfer Kit according to the manufacturer's instructions. The membrane was blocked with Bullet Blocking One (Nacalai USA 13779) then subjected to antibody reactions. After the secondary antibody reaction, Clarity Western ECL substrate (Bio-Rad) was applied and chemiluminescence images were captured using Bio-Rad ChemiDoc XRS.

#### E/L Repli-seq Library Preparation and Data Processing

Repli-seq was performed as previously described (Marchal et al. 2018). Briefly, cells were labeled with 100uM BrdU (Sigma Aldrich, B5002) for 2 hours and then fixed in 75% ethanol. Fixed cells were then sorted by FACS into early-S and late-S fractions based on propidium iodide staining of DNA. DNA was then purified from each fraction, sheared using Covaris ME220, and used to construct Illumina sequencing libraries with NEBNext Ultra DNA Library Prep Kit for Illumina (NEB, E7370). BrdU-labeled nascent DNA library fragments were then enriched by immunoprecipitation with anti-BrdU antibody (BD, 555627). IP products are then PCR amplified and indexed. A minimum of 10M reads/read pairs per library were targeted for each library. Raw sequencing data were first quality and adaptor trimmed using *cutadapt* from Trim Galore (Martin 2011) and then aligned to the mm10 reference genome with *bowtie2* (Langmead and Salzberg 2012). Aligned reads were then quality filtered and duplicate reads removed using *samtools* (Li et al. 2009). Processed aligned reads were then counted in 50kb windows across the entire genome for both early- and late-fraction using *bedtools* (Quinlan and Hall 2010) and then a log2 ratio of early-to-late read counts for each window was calculated to generate raw timing files. Raw E/L data was then scaled with R (Team 2021) and quantile normalized by dcHiC (Chakraborty et al. 2022). The change in RT ($\Delta_{RT}$) was calculated by subtracting the DMSO RT from the dTAG RT of combined replicates. For browser views and track-style visualizations, RT values or their delta were loess smoothed per chromosome in 300kb windows (Turner et al. 2023).

Micro-C data analysis

Raw sequencing data (fastq files) under GSE163729 were downloaded from GEO/SRA. The HiCPro (v.3.10) (Servant et al. 2015) analysis pipeline was used to obtain the contact matrices and valid pairs used in downstream analysis. Paired-end Micro-C reads were mapped with Bowtie2 (v.2.4.4) (Langmead and Salzberg 2012) to the *Mus Musculus* reference genome mm10 in *very-sensitive-local* mode. Pairs are then filtered to discard those marked as PCR duplicates, multiple hits, dangling end, self-circle or had low mapping quality scores. For verification, processed .hic files were also obtained from GEO and analyzed in parallel.

Compartment analysis

We initially utilized the preprocessed .hic files from GEO and performed compartment analysis using Juicer tools. Upon our inability to reproduce published results for chromosomes 11 and 17, we reprocessed the raw reads of Micro- C data and employed dcHiC for compartment calling at 100kb and 200kb resolution. Briefly, sparse matrices in HiCPro format were inputted into dcHiC (Chakraborty et al. 2022) and the first and second PCs for each chromosome are computed. Then, dcHiC utility tools were used to select the best PC for each chromosome. This PC choice and orientation of the PC sign was confirmed by calculating correlation with laminB signal (Peric-Hupkes et al. 2010).
